# Autologous mitochondria transport via transzonal filopodia rejuvenates aged oocytes by UC-MSCs derived granulosa cells-oocyte aggregation

**DOI:** 10.1101/2021.10.30.466630

**Authors:** Shuang Tang, Nannan Yang, Mingxi Yu, Shuo Wang, Xiangdong Hu, Heliang Ni, Wenyang Cai

**Author notes:** Corresponding author: Shuang Tang.; Address: Shenyang Agricultural University, 120 Dongling Road, Shenhe District, Shenyang, Liaoning 110866, China.

## Abstract

Mitochondria transfer can rescue oocyte aging-related infertility. However, heterologous techniques are suspended due to heteroplasmy. Regarding autologous approaches, the donor source and manipulating procedures require further optimization. Here we propose a strategy using umbilical cord mesenchymal stem cells (UC-MSCs) as mitochondria donor cells and employing intercellular mitochondria transport as the transfer method. We cryopreserved UC-MSCs of the female pup. When the female aged, its UC-MSCs were induced into granulosa cells (iGCs). The zona-weakened GV oocytes were aggregated with autologous iGCs into iGC-oocyte complexes. After cultivation in GDF9-containing media, mitochondria migrated from iGCs into the GV oocyte via transzonal filopodia. The maturation rate, quality, and developmental potential of these oocytes were substantially increased. Furthermore, the birth rate after embryo transfer has been improved. This approach utilized noninvasive procedures to collect mitochondria donor cells and optimized mitochondria transfer manipulations, so may represent a promising advance towards the improvement of aging-related infertility.

## Introduction

Many women of advanced reproductive age are suffering from infertility because the quality of their oocytes decreases with age ^1,2^. Mitochondria supply energy for meiosis and preimplantation embryo development ^3,4^. The insufficient quantity and dysfunction of mitochondria cause the quality decline of aged oocytes ^1,2^. Transfer of partial ooplasm ^5,6^, germinal vesicle (GV) ^7^, spindle ^8^, polar body ^9^, pronuclei ^10^, or nuclear genome ^11^ has been tested in animals and humans to replenish oocytes with healthy mitochondria. Although live birth of babies has been achieved, these heterologous techniques are not recommended due to the risk of heteroplasmy ^1^. Microinjection of autologous mitochondria from cumulus granulosa cells has been shown to improve oocyte quality and infertility in aged women ^12^. Nevertheless, it is worth noting that these granulosa cells also age with maternal aging ^13^. Transfer of autologous mitochondria from oogonial stem cells (OSCs) has also been raised to rejuvenate oocytes ^14^, while the existence of OSCs is still highly controversial ^15,16^. Moreover, highly invasive procedures are used to harvest OSCs, which increase the patient’s pains ^14,17^. Therefore, finding of new mitochondria donor source and optimization of manipulating procedures of autologous techniques are needed to solve these issues.

Previous studies illustrate that pluripotent stem cells contain undifferentiated, round, and globular immature mitochondria, similar to those in oocytes ^1,18,19^. Mesenchymal stem cells (MSCs) are attractive cells for their capacity of proliferation and multilineage differentiation ^20^. MSCs can differentiate into granulosa cells *in vivo* or *in vitro* ^21,22^. Moreover, MSCs rescue ovarian aging after differentiation into granulosa cells ^22^. The umbilical cord is a good source of MSCs (UC-MSCs) since it is younger than adult tissues. Moreover, UC-MSCs are collected using noninvasive procedures ^20^. Studies have demonstrated that UC-MSCs can be induced into oocyte-like cells in animals and humans ^23-25^. Thus, we speculated that UC-MSCs might be an ideal autologous mitochondria source for oocyte rejuvenation.

Mitochondria transport between neighboring cells was first documented in 2006 ^26^. Hereafter, studies showed that mitochondria migrate intercellularly via actin-based filopodia, termed tunneling nanotubes (TNTs) ^27,28^. In the investigations of germline-soma communication, Macaulay et al. and El-Hayek et al. reported that growth-differentiation factor 9 (GDF9) secreted by the oocyte induces the surrounding granulosa cells to generate specialized filopodia, termed transzonal projections (TZPs) in this case, which penetrate the zona pellucida and permit essential communication ^29,30^. An early electron microscopy study has identified a wide spectrum of mitochondrial sizes in mouse oocytes ranging from as small as 0.2 μm to as large as 0.6 μm or more ^31^. Although no organelle transport through TZPs has been reported so far, Macaulay et al. indicated that the diameter of TZPs can reach 2 μm ^30^, which is sufficient for mitochondria transport.

In this study, we examined whether UC-MSCs are an ideal source of autologous mitochondria. Then we tested the feasibility of a simple strategy for oocyte rejuvenation. By aggregating the zona-weakened GV oocyte with the granulosa cells induced from autologous UC-MSCs (iGCs), younger healthy mitochondria migrated into the oocyte through transzonal filopodia, which rejuvenated the aged oocyte and improved aging-related infertility.

## Results

### Isolation and characterization of UC-MSCs

We obtained pups using Cesarean section and isolated UC-MSCs from female pups. Primary cells could migrate out from umbilical cord explants after 1∼2 days and grow to 70∼80% confluence after about 9∼12 days of culture. The cells exhibited triangular, spindle-shaped morphology similar to fibroblasts (Fig. 1a). These cells could proliferate up to 40 passages (Fig. 1b). The doubling times of three randomly selected UC-MSCs for testing were 17.8 h, 19.1 h, and 18.8 h. Their characteristic MSC phenotypes were maintained after several passages, which could be determined by the expression of MSC marker genes including *Nanog, Oct4, Sox2, Rex1*, and *Tert* (Fig. 1c). Furthermore, immunofluorescent staining showed the cells were positive for OCT4, NANOG, SOX2, and KLF4 (Fig. 1d). The immunophenotypic characterization of the cells was analyzed by using flow cytometry. The data showed that the isolated cells were positive for MSC markers CD90 and CD105, yet negative for hematopoietic markers CD19 and CD45 (Fig. 1e). The isolated cells were cryopreserved for providing autologous UC-MSCs when the corresponding female aged.

**Figure 1.**
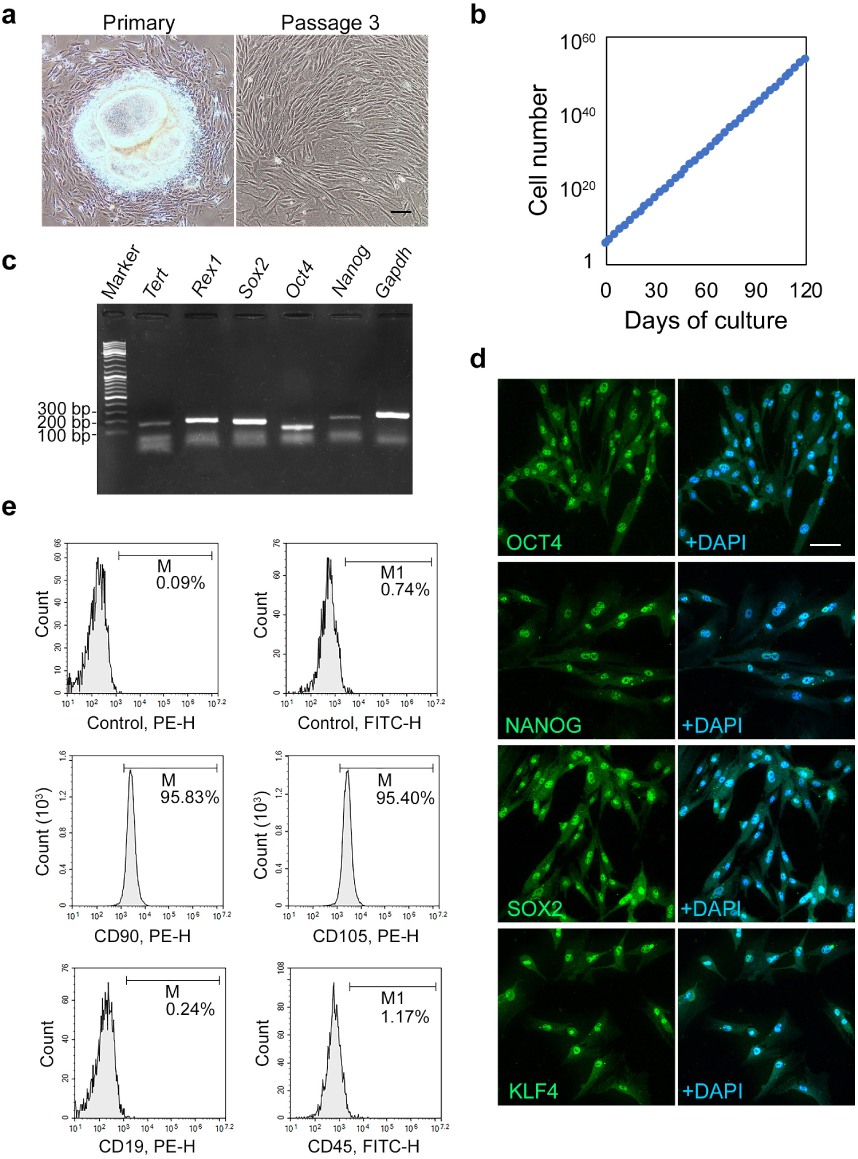
Characterization of UC-MSCs. **a**, Morphology of UC-MSCs. Scale bar = 100 μm. **b**, The growth curve of UC-MSCs. 2 × 10^5^ cells were passaged every 3 days into each well of 6-well plates. **c**, UC-MSCs were positive for *Nanog, Oct4, Sox2, Rex1*, and *Tert* analyzed by RT-PCR. *Gapdh* was used as a loading control. **d**, UC-MSCs were positive for OCT4, NANOG, SOX2, and KLF4 analyzed by immunofluorescence. Scale bar = 100 μm. **e**, UC-MSCs were positive for CD90 and CD105 and negative for CD19 and CD45 analyzed by flow cytometry.

### UC-MSCs as an ideal source of autologous mitochondria

To investigate whether UC-MSCs are ideal donor cells, aged GV oocytes were injected with mitochondria isolated from autologous UC-MSCs. The injected oocytes were reaggregated with their cumulus cells and subjected to *in vitro* maturation and fertilization. First, we checked the first polar body extrusion. The maturation of mitochondria-injected oocytes was effectively improved in comparison with buffer-injected controls (Fig. 2a). We then determined the oocyte quality by examining the ATP content, intracellular reactive oxygen species (ROS) levels, and spindle organization in mature oocytes. The average ATP content was increased to 775.1 ± 15.5 fmol/oocyte in the mitochondria-injected group compared with 371.9 ± 11.7 fmol/oocyte in the control group (Fig. 2b). In contrast with the controls, mitochondria-injected oocytes showed a significant decrease in intracellular ROS levels (Fig. 2c, d). The percentage of abnormal spindles with disordered microtubule structure and/or chromosome misalignment was markedly reduced in the mitochondria-injected group (Fig. 2e, f). These results demonstrate that the injection of autologous mitochondria from UC-MSCs could substantially improve the quality of aged oocytes.

**Figure 2.**
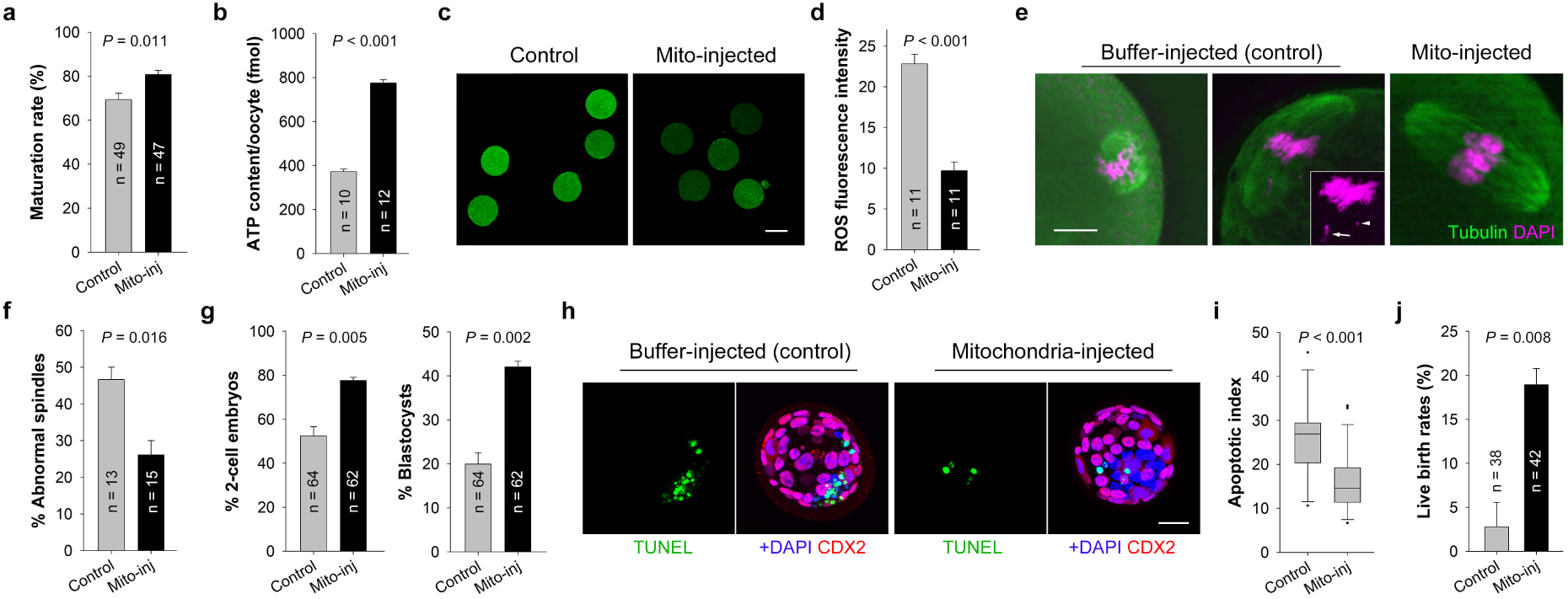
Microinjection of autologous mitochondria from UC-MSCs improves the quality of aged oocytes. **a**, The *in vitro* maturation rates of buffer-injected (control) and mitochondria-injected (Mito-inj) oocytes. **b**, The ATP content per oocyte of the control and mitochondria-injected groups. **c**, Representative confocal images of intracellular ROS in control and mitochondria-injected oocytes. Scale bars= 50 μm. **d**, Quantitative analysis of ROS fluorescence intensity. **e**, Representative images of spindles in control and mitochondria-injected oocytes. Scale bar = 10 μm. The middle panel shows an abnormal spindle having nearly normal microtubule structure yet with misaligned and detached chromosomes (please refer to Criteria of spindle scoring in Methods). The inset shows detached (arrow) and fragmented (arrowhead) chromosomes. **f**, The percentages of abnormal spindles in the control and mitochondria-injected groups. **g**, The *in vitro* developmental rates of 2-cell embryos and blastocysts in the control and mitochondria-injected groups. **h**, Representative images of blastocysts examined by TUNEL assay. Scale bar = 20 μm. **i**, The box plots of apoptotic index for blastocysts from the control (n = 13) and mitochondria-injected (n = 26) groups. Dots indicates outliers. **j**, Live birth rates after transferring of 2-cell embryos from the control or mitochondria-injected group into surrogate mothers. Data were analyzed by Student’s *t*-test. Each bar denotes mean ± SEM.

The developmental potential of the injected oocytes was evaluated by monitoring the *in vitro* embryo development, the apoptotic incidence in blastocysts, and birth rates after embryo transfer. The rates of 2-cell embryo development and blastocyst formation were significantly increased from fertilized oocytes in the mitochondria-injected group (Fig. 2g). Moreover, reduced apoptotic cells were observed in blastocysts from the mitochondria-injected group. (Fig. 2h, i). In embryo transfer experiments, the 2-cell embryos were transferred into oviducts of pseudopregnant females. Eight live births were resulted after transferring 42 embryos into 3 recipients in the mitochondria-injected group, while only one pup was obtained after transferring 38 embryos into 3 recipients in the control group (Fig. 2j). Therefore, we could see the obvious effects of autologous mitochondria from UC-MSCs on fertility improvement in aged female mice.

Collectively, the above evidence suggests that UC-MSCs are an ideal donor source of autologous mitochondria.

### Induction of UC-MSCs into granulosa cells (iGCs)

For induction of iGCs, UC-MSCs were cocultured with granulosa cells in conditioned media. With the induction, the morphology of UC-MSCs gradually changed into a granulosa cell-like shape (Fig. 3a). RT-PCR analysis revealed that these cells expressed granulosa cell markers, including *Foxl2, Cyp19a1, Amhr2, Amh*, and *Fshr* (Fig. 3b). Immunofluorescent staining showed these cells were positive for FOXL2, AMH, AMHR2, and FSHR (Fig. 3c). We then generated iGCs from UC-MSCs transfected with *COX8*-tdTurboRFP, so the mitochondria in these cells were labeled with the red fluorescent protein (RFP) (Fig. S1). After overnight culture in GDF9-containing media, the iGCs could respond to GDF9 and form extended filopodia. Between neighboring iGCs, mitochondria transport was observed via intercellular filopodia (Fig. 3d). To examine whether iGCs could inhibit meiotic resumption of GV oocytes in GDF9-containing media, we used different cells to construct cell-oocyte complexes. iGCs could maintain meiotic arrest of GV oocytes in at least 5 days of culture, which was similar to granulosa cells. In contrast, GV oocytes aggregated with UC-MSCs quickly resumed meiotic progression (Fig. 3e). These results demonstrate that UC-MSCs can be induced into functional granulosa cells.

**Figure 3.**
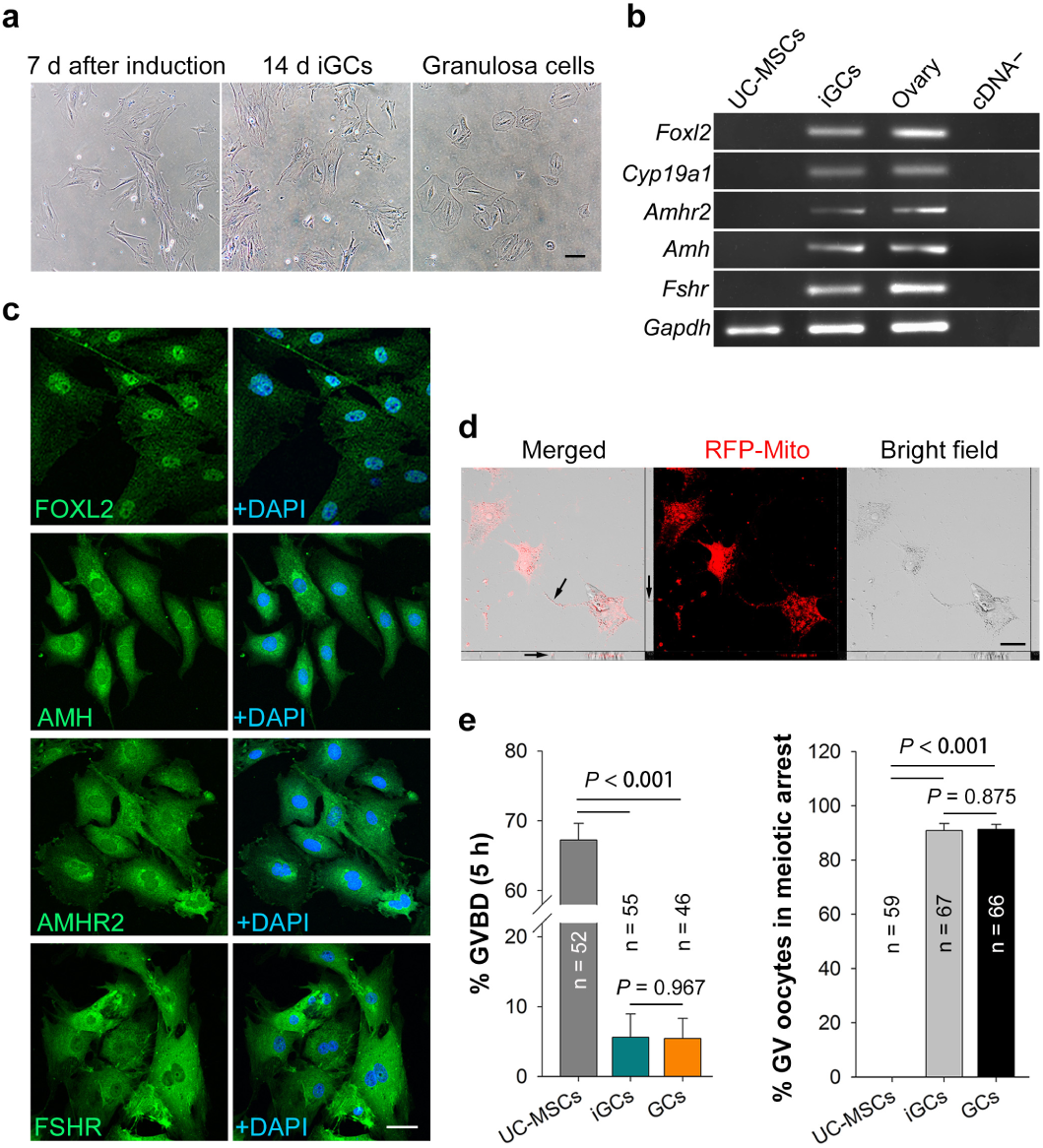
Generation of iGCs from UC-MSCs. **a**, Morphology of UC-MSCs after 7 days of induction, iGCs (14 days after induction), and granulosa cells. **b**, RT-PCR analysis of granulosa cell marker genes in UC-MSCs, iGCs, and ovary. *Gapdh* was used as a loading control. **c**, Immunofluorescence showing iGCs were positive for FOXL2, AMH, AMHR2, and FSHR. **d**, Mitochondria transport via intercellular filopodia generated between neighboring iGCs in response to GDF9. RFP-Mito means RFP-labeled mitochondria. Arrows indicate a mitochondrion in the intercellular filopodium. **e**, Monitoring of GVBD (germinal vesicle breakdown) and GV oocyte maintenance showed that iGCs had the same function as granulosa cells (GCs) to maintain meiotic arrest of GV oocytes in GDF9-containing media. The data were analyzed by one-way ANOVA followed by a Holm-Sidak test. Each bar represents mean ± SEM. All scale bars = 100 μm.

### Zona weakening facilitates mitochondria transport in iGC-oocyte complexes

We have seen that iGCs can form intercellular filopodia in response to GDF9. Next, we wanted to know the conditions under which mitochondria transport may occur after aggregating aged GV oocytes with iGCs. We used mitochondria-labeled iGCs to construct iGC-oocyte complexes for monitoring mitochondria transport. After 3 days of cultivation in GDF9-containing media, some of the iGCs were removed by repeated aspiration and iGC-oocyte complexes were observed under a confocal microscope. In iGC-zona-intact oocyte aggregates, we did not, however, observe overt mitochondria transport into the GV oocytes (Fig. 4a, b).

**Figure 4.**
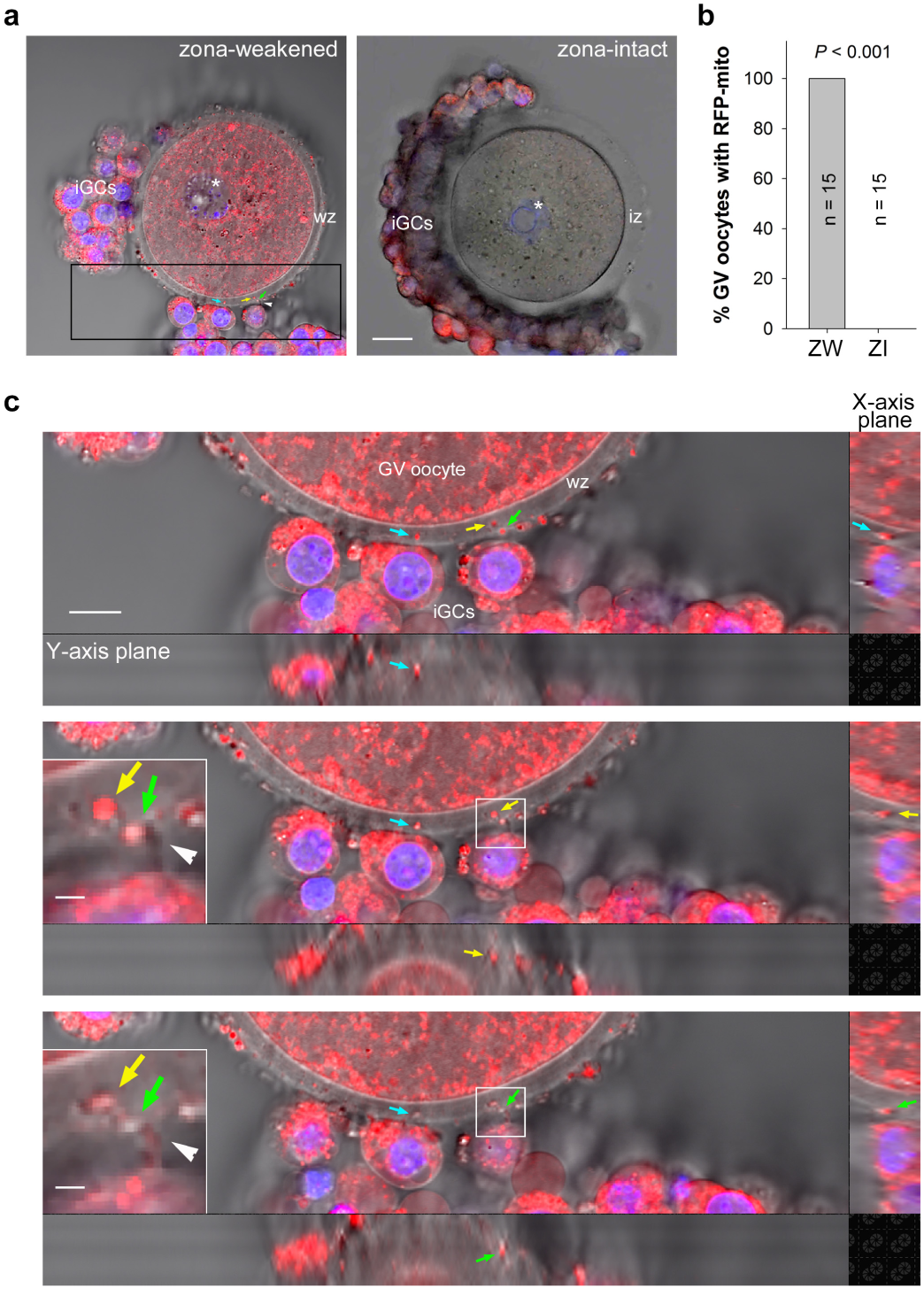
Mitochondria transport via transzonal filopodia in iGC-oocyte complex. **a**, Representative images of an iGC-zona-weakened GV oocyte complex and an aggregate constructed by iGCs and a zona-intact GV oocyte. The mitochondria originating from iGCs had been labeled with RFP and can be identified by red fluorescence. Blue fluorescence represents nuclei stained with Hoechst 33342. Asterisks mark the germinal vesicle. “wz” means weakened zona and “iz” means intact zona. Note their different thickness. The arrows with different colors indicate the different mitochondrion traversing the zona pellucida. The arrowhead indicates a transzonal filopodium. Scale bar = 20 μm. **b**, The percentages of GV oocytes carrying iGC mitochondria in the zona-weakened (ZW) and zona-intact (ZI) groups. Data were analyzed by Student’s *t*-test. Each bar represents mean ± SEM. **c**, Three consecutive Z-stack images of the boxed region in panel **a** show mitochondria traversing the zona pellucida via transzonal filopodia. Scale bar = 10 μm. The insets show two mitochondria were migrating into the oocyte one after another through a transzonal filopodium (arrowhead) budding from an iGC. Scale bar = 1 μm.

Since the zona thickness of growing oocytes may reach 6.2 μm ^32^, it was reasonable to speculate that zona pellucida might obstruct organelle transport between iGCs and oocytes. To test this, we placed zona-free GV oocytes on iGCs in GDF9-containing media. As can be seen, RFP-labeled mitochondria gradually appeared in the oocyte over time (Fig. S2). Nonetheless, the absence of zona pellucida leads to polyspermy, and declines in oocyte developmental potential and embryo quality ^32^. Therefore, it is impracticable to completely remove the zona pellucida in clinical practice.

Therefore, we tried to weaken the zona pellucida rather than remove it. After weakening of the zona by using Tyrode’s solution, mitochondria transport occurred in the aggregated complexes (Fig. 4). RFP-labeled mitochondria were clearly visible in the ooplasm (Fig. 4a, c), indicating that mitochondria had migrated from iGCs into the GV oocyte. Figure 4c showed enlarged images of the iGC-zona-oocyte boundary area. It can be seen that mitochondria were traversing the zona pellucida via the filopodia protruding from iGCs. These results demonstrate that zona weakening can facilitate mitochondria transport from iGCs into the GV oocyte via transzonal filopodia. To further verify this finding, we punched several pores on the zona with a microcapillary and then constructed iGC-oocyte complexes. The wider filopodia could be formed between iGCs and the oocyte, through which mitochondria were migrating (Fig. S3). This observation confirms the above conclusion.

### Rejuvenation of aged oocytes by mitochondria from iGCs

We next investigated the effects of mitochondria transport on the quality and developmental potential of aged oocytes. Zona-weakened GV oocytes were aggregated with autologous iGCs, and these complexes were cultured in GDF9-containing media for 3 days. First, mitochondrial ultrastructure in GV oocytes was analyzed by using TEM. As shown in Figure 5a, the treated GV oocytes (from the iGC-zona-weakened GV oocyte complex group) mainly contained normal mitochondria which displayed well-aligned outer and inner membranes and well-defined intermembrane space. In contrast, the control oocytes had many abnormal mitochondria with ultrastructural features including the mitochondrial vacuole and/or a narrowed intermembrane space. Quantitative analysis of different ooplasm regions of several oocytes showed that the proportion of abnormal mitochondria in treated oocytes was considerably lower than that in controls (Fig. 5b). Then, we examined the ATP content and intracellular ROS levels in GV oocytes. ATP measurements revealed that mitochondria from iGCs dramatically increased the ATP level in aged GV oocytes over that in controls (816.1 ± 13.3 fmol/oocyte vs. 393.6 ± 10.1 fmol/oocyte; Fig. 5c). The treated oocytes showed significantly reduced ROS levels compared with the controls (Fig. 5d, e). These results demonstrate that mitochondria transport by iGC-oocyte aggregation substantially improved the quality of aged GV oocytes.

**Figure 5.**
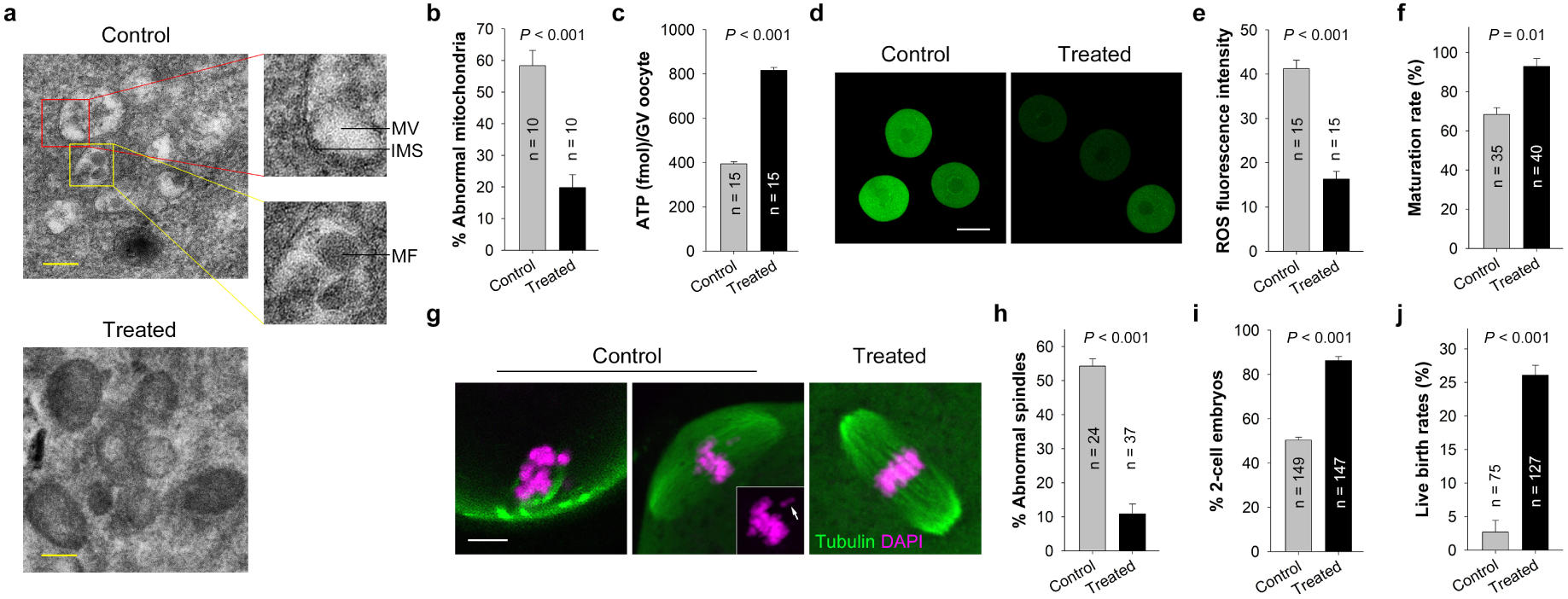
Rejuvenation of aged oocytes by mitochondria transport. **a**, Representative TEM micrographs of mitochondria from GV oocytes in the control and treated (iGC-zona-weakened GV oocyte complex) groups. Scale bar = 200 nm. IMS: intermembrane space, MV: mitochondrial vacuole, MF: myelin figures. **b**, The proportion of abnormal mitochondria in GV oocytes of the control and treated groups. **c**, The ATP content per GV oocyte of the control and treated groups. **d**, Representative images of intracellular ROS in GV oocytes of the control and treated groups. Scale bars = 50 μm. **e**, Quantitative analysis of ROS fluorescence intensity. **f**, The *in vitro* maturation rates of control and treated oocytes. **g**, Representative confocal images of spindles in mature oocytes from the control and treated groups. Scale bars = 10 μm. The middle panel shows an abnormal spindle having nearly normal microtubule structure yet with misaligned and detached chromosomes. The inset shows the chromosome (arrow) detached from the spindle equator. **h**, The percentages of abnormal spindles in the control and treated groups. **i**, The *in vitro* developmental rates of 2-cell embryos in the control and treated groups. **j**. Live birth rates after transferring of 2-cell embryos from the control or treated group into surrogate mothers. Data were analyzed by Student’s *t*-test. Each bar represents mean ± SEM.

The iGC-oocyte complexes were subjected to *in vitro* maturation and fertilization. The maturation rate of treated oocytes was markedly higher than that of controls (Fig. 5f). A striking reduction was also noted in the percentage of abnormal spindles in mature oocytes from the treated group (Fig. 5g, h). The in *vitro* developmental rate of 2-cell embryos was significantly increased from fertilized oocytes in the treated group (Fig. 5i). In embryo transfer experiments, the 2-cell embryos were transferred into oviducts of pseudopregnant females. Thirty-three births were resulted after transferring 127 embryos into 12 recipients in the treated group, while only two pups were obtained after transferring 75 embryos into 6 recipients in the control group (Fig. 5j). These results demonstrate that, by aggregating the zona-weakened GV oocyte with autologous iGCs, younger healthy mitochondria from iGCs rejuvenated the aged oocyte and improved aging-related infertility.

## Discussion

Here we propose a simple strategy for oocyte rejuvenation. First, we induced autologous UC-MSCs into granulosa cells (iGCs). Then, we aggregated the zona-weakened GV oocyte with iGCs into a complex. Next, we cultured the complex in GDF9-containing media. By this means, mitochondria migrated into the oocyte via transzonal filopodia budding from iGCs and rejuvenated the aged oocyte (Fig. 6). Our results demonstrate that mitochondria transport can occur between the surrounding granulosa cells and the oocyte, which is critical for the oocyte quality. This conclusion can be mutually corroborated with the previous findings that the mitochondrial DNA copy number of cumulus granulosa cells is strongly correlated with the oocyte quality in clinical research and animal studies ^33-36^. Evidence has been presented that the oocyte directly connects to granulosa cells by fusing with the cell membrane during follicle development ^37^. In fact, scientists have already hypothesized that transzonal filopodia may serve the function of organelle transport from the cumulus cell body to the oocyte, yet there was no evidence ^30,38^. Our finding provides direct evidence for this idea.

**Figure 6.**
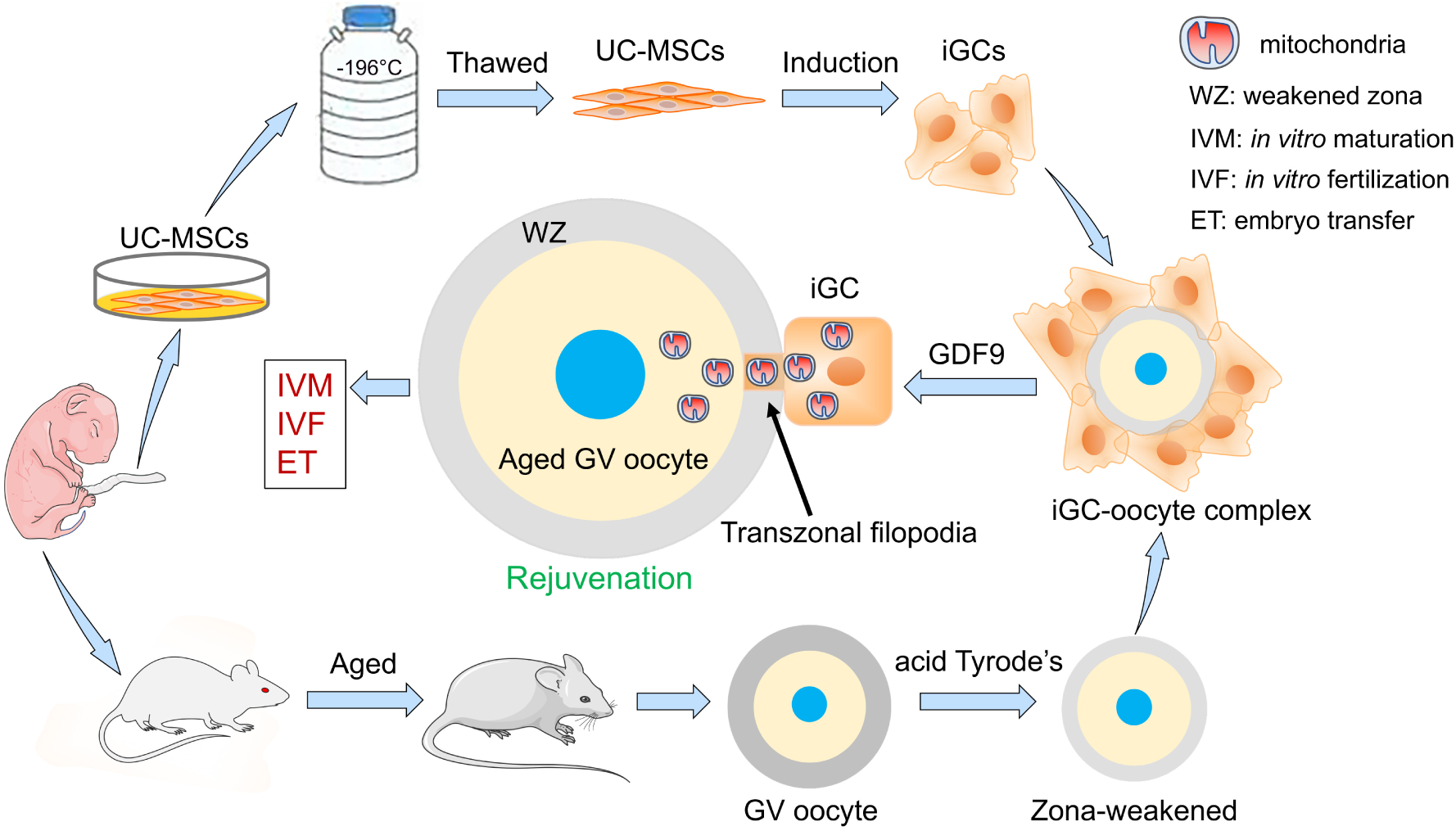
The strategy of oocyte rejuvenation by iGC-oocyte aggregation. UC-MSCs were isolated from the umbilical cord of female pups delivered by Cesarean section on day 18 or day 19 and cryopreserved in liquid nitrogen. When the female grew elder, its UC-MSCs were induced into iGCs (induced granulosa cells). GV oocytes were collected from the aged mouse and treated to weaken the zona pellucida. The zona-weakened GV oocytes were aggregated with autologous iGCs into iGC-oocyte complexes. After cultivation in GDF9-containing media, younger healthy mitochondria migrated into the GV oocyte via transzonal filopodia budding from iGCs. The iGC-oocyte complexes were subjected to *in vitro* maturation and fertilization. The cleaved 2-cell embryos were selected for embryo transfer.

Of note, the occurrence of such mitochondria transport requires certain conditions. Using zona-intact GV oocytes, we could not observe overt mitochondria transport (Fig. 4a, b). In previous cell experiments, as long as TNTs, a type of specialized filopodia structure, are formed between neighboring cells, mitochondria transport can take place ^26-28^. Furthermore, videomicroscopy has revealed the presence of vesicle-like structures moving along traversing filopodia connecting mural trophectoderm with inner cell mass in mouse blastocysts ^39^, implying that organelle transport via intercellular filopodia might be a normal physiological activity in the embryo. In the early stage of follicular development, the oocyte and the surrounding granulosa cells are very tightly connected as a syncytium with close communication ^32,37,40^. As the oocyte grows, its zona increases from about 1.6 to 6.2 μm in width ^32^. From these facts, we speculated that the pores in the zona are gradually narrowing with oocyte growth so that the width of transzonal filopodia becomes insufficient for mitochondria to pass through. In our experiments, we found that, after weakening of the zona pellucida with Tyrode’s solution, efficient mitochondria transport was observed between iGCs and the oocyte (Fig. 4). We also found that zona punching can facilitate the formation of wider transzonal filopodia for mitochondria transport (Fig. S3). These findings indicate that zona weakening is a critical procedure in the proposed strategy.

Compared with the cell types in prior studies, we suggested that UC-MSCs may be an optimal supplying source of autologous mitochondria. A clinical trial has proposed that using autologous cumulus granulosa cells as mitochondria donor cells can rescue aged oocytes and obtain live births ^12^. But the problem is that these somatic cells also age with oocyte aging ^13^. Studies have shown that the qualities of oocytes and granulosa cells are positively correlated ^33-36^. Obviously, aged granulosa cells are not a good candidate. Adult stem cells are also used as mitochondria donor cells. It has been reported that mitochondria from autologous adipose-derived stem cells (ADSCs) have normal morphology and can improve oocyte quality and infertility in aged mice ^19^. Nevertheless, another investigation using mitochondria from autologous ADSCs has reached a conflicting conclusion ^13^. Therefore, adult ADSCs are not necessarily a suitable mitochondria source. Unlike these cells, UC-MSCs are fetal stem cells isolated from birth-associated tissue, younger than adult stem cells ^20^. Moreover, studies have demonstrated that UC-MSCs can be induced into oocyte-like cells ^23-25^. Our results demonstrated a clear improvement in either oocyte quality or birth rates when mitochondria from UC-MSCs or iGCs were provided (Fig. 2, Fig. 5). The previous finding that MSCs rescue ovarian aging by differentiation into granulosa cells can also be supporting evidence to our conclusion ^22^.

Another exciting candidate source of mitochondria is OSCs ^14^. Regrettably, their existence is widely questioned. A joint study conducted by multiple independent laboratories provides evidence denying the existence of DDX4-expressing functional OSCs in adult human and mouse ovaries ^16^. The single-cell analysis also demonstrates no OSCs in the human ovarian cortex ^15^. Furthermore, the putative OSCs are collected from cortical tissue dissected by laparoscopic ovarian biopsy ^14,17^. The manipulations comprise highly invasive procedures and therefore have limited applicability ^1^. In contrast, our proposed strategy employs noninvasive procedures to obtain mitochondria without involving the ovarian tissue. The procedures only need to isolate and cryopreserve UC-MSCs at the birth of female babies and then these cells can be used after the females grow older ^20^. ADSCs collection also involves invasive procedures ^13,19^. There is no harm to the mother, which is also a great advantage of UC-MSCs.

Additionally, in comparison with previous approaches, our proposed strategy does little harm to the oocyte. Microinjection or nuclear transfer used in previous strategies gives rise to mechanical damages to the oocyte membrane and cytoskeleton. Moreover, they might also disrupt the spindle apparatus and cause chromosomal abnormalities in the developing embryo ^1,41-43^. The iGC-oocyte aggregation circumvents the micromanipulation of oocytes, thus avoiding these damages. In previous strategies, chemical activation has been used to activate oocytes ^1,7^. Nevertheless, artificial calcium waves may have effects on downstream molecular events ^44^. Electrofusion used in ooplasm transfer or spindle transfer might induce oocyte premature activation and thereby cause high aneuploidy rates ^1,6,45,46^. In contrast, our strategy excludes these manipulations. iGC-oocyte complexes are similar to cumulus-oocyte complexes. The embryos for transfer can be obtained after iGC-oocyte complexes undergo *in vitro* maturation and fertilization, avoiding the potential adverse effects of artificial calcium waves or electric pulses. Overall, the proposed strategy may be more conducive to embryonic development and offspring health.

The timing of mitochondria transport may be another determinant of the efficacy of rejuvenation strategies. Except for GV transfer, in some of the prior strategies, mitochondria are supplied to MII oocytes ^1,2^. However, the highest risk of aging-related misalignments is associated with the first meiotic division, since mitochondrial ATP synthesis provides energy for organization of microtubules during meiotic spindle assembly ^1,47^. In this study, we found a significant increase of ATP content in GV oocytes from the treated group compared with those from the control group, which substantially decreased the probability of spindle defects and chromosomal misalignments (Fig. 5c, g, h). In our proposed strategy, mitochondria are supplied to the GV oocyte, which can as early as possible reduce meiotic abnormalities caused by energy insufficiency.

In addition to mitochondria transport, aggregation facilitates substance communication (*e*.*g*. RNAs, nutrients, substrates for antioxidant synthesis) between oocytes and iGCs ^29,30,48,49^, which might also contribute to improving oocyte quality.

Our findings show that mitochondria can migrate into the oocyte through transzonal filopodia budding from iGCs, but did not exclude the possibility of other transport manners such as microvesicles, exosomes, or expanded gap junctions ^40,50^. This requires further exploration to uncover all transport mechanisms. In addition, whether the oocyte can select high-quality mitochondria during the transport process is also an interesting question which needs to be answered in future investigations. Nevertheless, we believe this new design of autologous approach may represent a promising advance towards the improvement of aging-related infertility.

## Acknowledgments

We are grateful to Dr. Xuyuan Yang in Wanlei Biotechnology Company for her assistance in flow cytometry. This study is funded by the Natural Science Fund Project, Department of Science and Technology of Liaoning Province (Grant number 2019-MS-282), Liaoning Science Planning Project, the Educational Department of Liaoning Province (Grant number LSNJC202009), and the National Natural Science Foundation of China (Grant number 31301201).

## Author contributions

**Shuang Tang:** Conceptualization, Methodology, Investigation, Validation, Formal analysis, Writing - Original draft, Writing - Review & Editing, Project administration, Funding acquisition. **Nannan Yang:** Investigation, Validation, Formal analysis, Data Curation. **Mingxi Yu:** Investigation, Formal analysis, Writing - Review & Editing. **Shuo Wang:** Methodology, Investigation. **Xiangdong Hu:** Investigation, Validation, Formal analysis. **Heliang Ni:** Investigation, Validation, Formal analysis. **Wenyang Cai:** Investigation, Formal analysis.

## Abbreviations

ADSCs: adipose-derived stem cells
BSA: bovine serum albumin
COCs: cumulus-oocyte complexes
DAPI: 4’,6-diamidino-2-phenylindole
EGF: epithelial growth factor
GDF9: growth-differentiation factor 9
FBS: fetal bovine serum
FSH: follicle-stimulating hormone
GV: germinal vesicle
GVBD: germinal vesicle breakdown
iGCs: induced granulosa cells from UC-MSCs
ITS: insulin–transferrin–selenium
IVF: *in vitro* fertilization
IVM: *in vitro* maturation
MSCs: mesenchymal stem cells
OSCs: oogonial stem cells
PMSG: pregnant mare serum gonadotropin
RFP: red fluorescent protein
ROS: reactive oxygen species
TEM: transmission electron microscope
TNTs: tunneling nanotubes
TZPs: transzonal projections
UC-MSCs: umbilical cord mesenchymal stem cells

---

**Figure S1.**
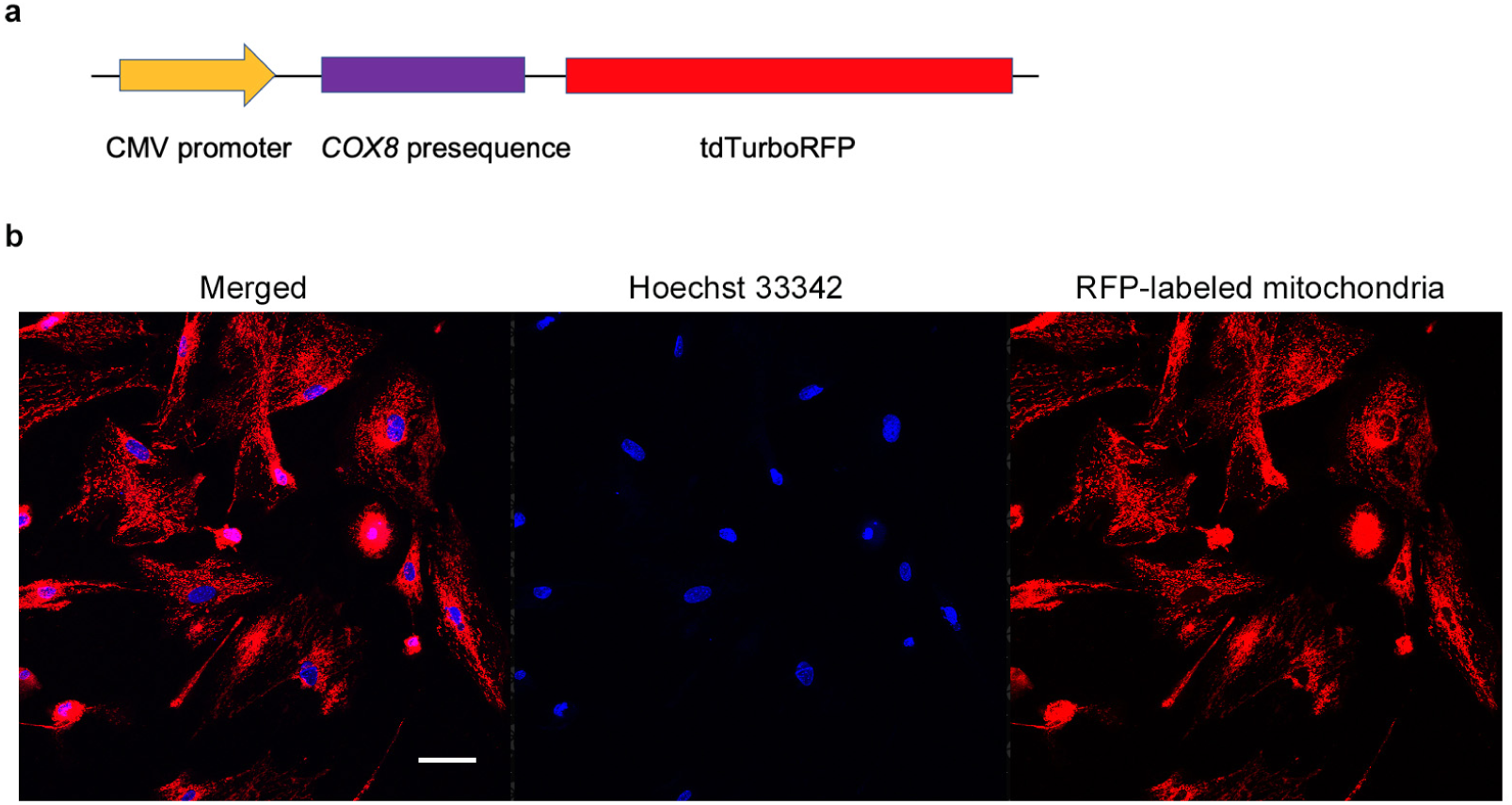
UC-MSCs transfection. **a**, The diagram of *COX8*-tdTurboRFP cassette. **b**, The RFP-labeled mitochondria in a strain of transgenic positive UC-MSCs. Scale bar = 100 μm.

**Figure S2.**
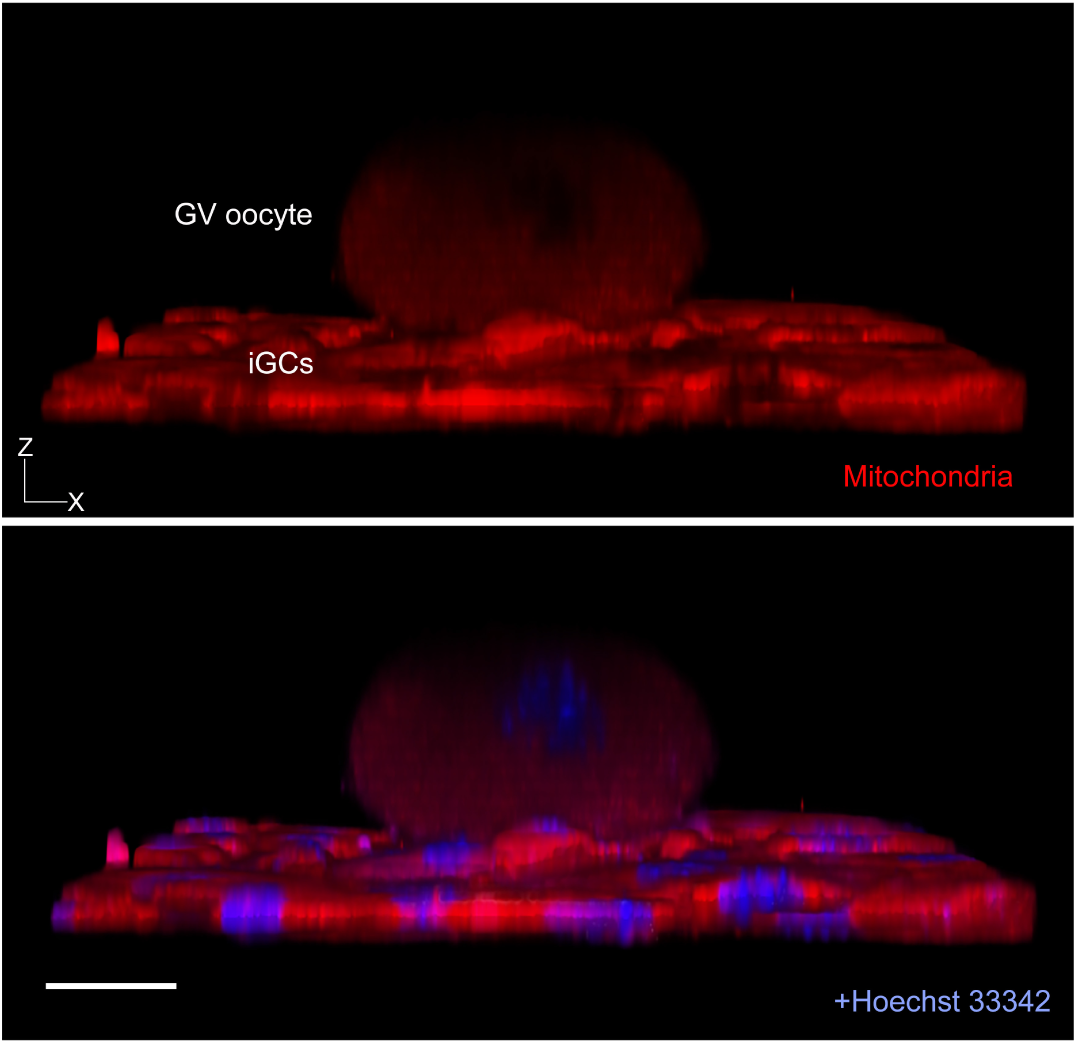
Mitochondria transport between zona-free GV oocytes and iGCs. The X-Z view of a volume rendered image from Z-stack data. Zona-free GV oocytes were placed on mitochondria-labeled iGCs in GDF9-containing media. Mitochondria transport into the oocyte was monitored at 6 h later. The mitochondria from iGCs were identified by red fluorescence. Scale bar = 20 μm.

**Figure S3.**
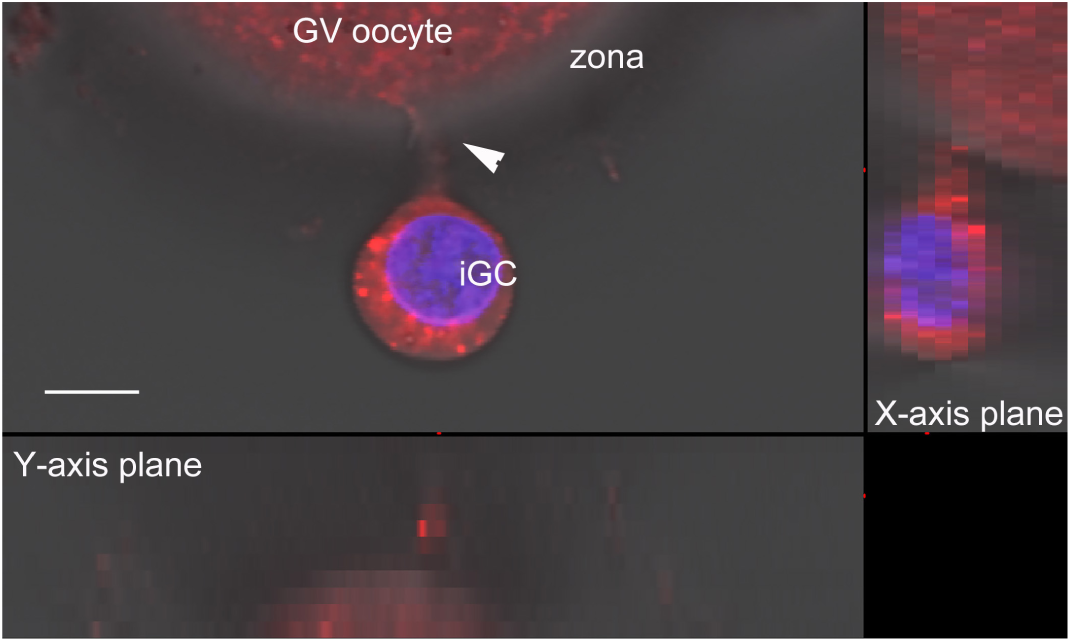
Zona punching facilitates the formation of wider transzonal filopodia for mitochondria transport. The zona-punched GV oocyte was aggregated with mitochondria-labeled iGCs. After cultivation in GDF9-containing media, some of the iGCs were removed by repeated aspiration and iGC-oocyte complexes were observed under a confocal microscope. The mitochondria originating from iGCs had been labeled with RFP and can be identified by red fluorescence. Blue fluorescence represents Hoechst 33342 staining. The arrowhead indicates a transzonal filopodium. Scale bar = 10 μm.

