## Supplementary material for "Autologous mitochondria transport via transzonal filopodia rejuvenates aged oocytes by UC-MSCs derived granulosa cells-oocyte aggregation": Methods

### Reagents

All chemicals were purchased from Sigma-Aldrich (China) LLC. (Shanghai, China) unless otherwise stated. DMEM-LG (10567), MEM-alpha (32571), HEPES-MEM (42360), Opti-MEM (51985034), and FBS (fetal bovine serum, 12664025) were from Gibco (Thermo Fisher Scientific (China) LLC., Shanghai, China). GDF9 (Mouse growth-differentiation factor 9, 739-G9-010) was purchased from R&D Systems (Minneapolis, MN). EGF (epithelial growth factor, 315-09) and ITS (insulin–transferrin–selenium, 00-102) were from PeproTech (Cranbury, NJ).

### Animals

ICR mice were purchased from Beijing Vital River Laboratory Animal Technology Co., Ltd. (Beijing, China). The mice were maintained with food and water *ad libitum* on a 12 h light/dark cycle under controlled temperature (23 to 25°C) in the SPF Lab Animal Center of Shenyang Agricultural University. All experiments were performed in compliance with the Guide for the Care and Use of Laboratory Animals and approved by the Institutional Animal Care and Use Committee of Shenyang Agricultural University.

### Cesarean section and fostering of the female pups

The 6 to 8-week-old female mice were caged with 12 to 24-week-old male mice to induce pregnancy (vaginal plug = day 1 of pregnancy). The pregnant female was sacrificed by cervical dislocation at 2200 h on day 18 or day 19 according to its pregnant behavior. The abdomen was disinfected with 75% ethanol and then laparotomized. The uterus was dissected and put onto sterile gauze with blunt forceps. Carefully cut open the uterine wall and took out the whole embryos with amnion and placenta. Each pup with its umbilical cord was dissected from the amniotic sac. Female pups were distinguished by observation of the anogenital distance under a stereoscope and numbered by toe-clipping. After numbering, the corresponding umbilical cord was cut off for UC-MSCs primary culture. The pups were then transferred to clean sterile gauze on a 37°C heating plate. Gently rolled the pups over for cleaning their bodies and kept them warm until they could automatically breathe. The pups were gently smeared with the foster mother's beddings. Then these pups were put into the cage to substitute the same number of the original ones. After weaning, these female mice were raised to 10~12 months old to provide aged GV oocytes for aggregation with its autologous UC-MSCs derived granulosa cells (iGCs).

**Establishment of umbilical cord mesenchymal stem cells (UC-MSCs)**

The individual numbered umbilical cord was rinsed twice in PBS, once in DMEM-LG supplemented with 10% FBS. Then, the umbilical cord was minced into small pieces with fine scissors. These explants were attached to the dish bottom and cultured in DMEM-LG plus 10% FBS in a humidified incubator at 37°C with 5% $CO_2$. The explants were removed after 2~3 days of cell migration. The media were refreshed every 3 days. The cells were passaged when reached 70~80% confluence. The cells of the fourth passage were cryopreserved in DMEM-LG containing 20% FBS and 10% dimethyl sulfoxide in liquid nitrogen for providing autologous UC-MSCs after the corresponding mouse grew elder.

**RT-PCR for marker genes**

Total RNA was extracted by using Trizol (Invitrogen, Thermo Fisher) and cDNA was synthesized with GoScrip Reversion Transcription System (Promega (Beijing) Biotech Co.,

Beijing, China). RT-PCR was performed using Premix *Taq* (Takara Bio (Dalian) Co., Ltd., Dalian, China). In the empty control, cDNA was substituted with ultrapure water. The program was as follows: 95°C × 5 min, followed by 30 cycles of 95°C × 30 s, 56°C × 30 s, 72°C × 30 s. Primers for RT-PCR were listed in Table S1. All primers have been tested using E13.5 whole embryo cDNA.

**Immunofluorescence**

The reagents for fixation, blocking and antibody dilution were purchased from Beyotime (Jiangsu, China). Cells were grown on poly-lysine-coated coverslips. Between two steps, cells or oocytes were washed three times in PBS plus 1 mg/mL polyvinylpyrrolidone-40 (PBS/PVP) for 5 min. Samples were fixed for 30 min at room temperature and permeabilized in 0.2% Triton X-100 in PBS for 4 min. Then they were blocked for 1 h at room temperature. Samples were incubated in primary antibodies overnight at 4 °C. Samples were then washed extensively and incubated in secondary antibodies for 20 min. The nuclei were counterstained with 1 μg/mL DAPI (4',6-diamidino-2-phenylindole) for 10 min. In negative control experiments, a subset of samples was stained with secondary antibodies and DAPI only. Samples were mounted on slides and observed under an A1+ laser confocal microscope (Nikon, Tokyo, Japan). Primary antibodies were listed in Table S2. Secondary antibodies (1: 800 dilutions) were from Jackson ImmunoResearch (West Grove, PA), listed in Table S3.

**Flow cytometry**

The surface antigen phenotyping of UC-MSCs was analyzed by flow cytometry. Single-cell suspensions were harvested by trypsinization and resuspended in PBS. The cells were incubated with FITC- or PE-conjugated antibodies (1: 160 dilutions) for 30 min at 4°C. PE-conjugated rat IgG2a and FITC-conjugated rat IgG2b were used as the isotype control. The detailed information on antibodies was listed in Table S4. After three times of washing with PBS, cells were analyzed with a NovoCyte flow cytometer (ACEA Biosciences, San Diego, CA). Data were analyzed with NovoExpress software. Single cells were determined by FSC-H/SSC-H. The Boundaries between “positive” and “negative” were defined by comparison with the isotype control. The gating strategy was shown in Figure S4.

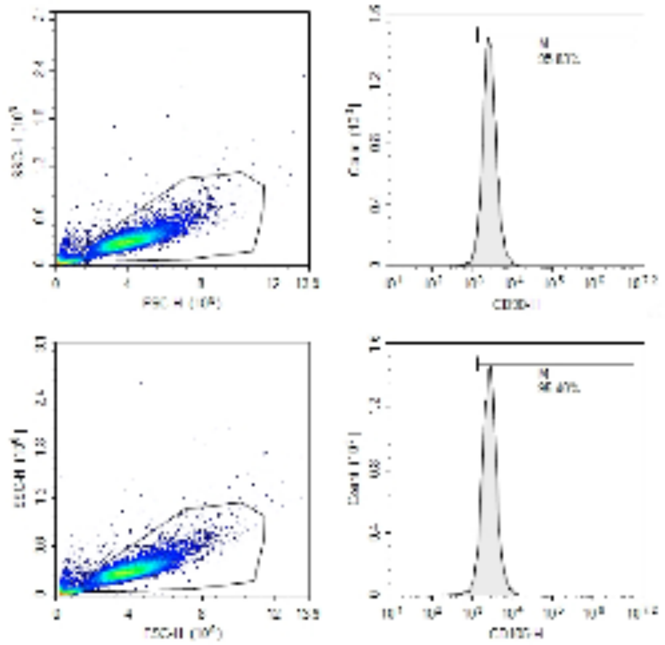

**Figure S4. Flow cytometry gating strategy.** Single cells determined by FSC-H/SSC-H. Cell debris was removed during the test (cell population in the lower left corner). The Boundaries between “positive” and “negative” were defined by comparison with the isotype control.

**Collection of GV oocytes**

Aged female (10~12 months old) was intraperitoneally injected with 10 IU pregnant mare serum gonadotropin (PMSG) (Ningbo Second Hormone Factory, Zhejiang, China). The mouse was sacrificed 46~48 h later and dissected ovaries were placed in HEPES-MEM supplemented with 1 mg/mL bovine serum albumin (BSA) and 0.5 IU/mL heparin. Antral follicles were punctured using a needle to release cumulus-oocyte complexes (COCs). GV oocytes were obtained by repeated aspiration with a fine-bore pipette for removing cumulus granulosa cells.

**Mitochondria isolation from UC-MSCs and microinjection into GV oocytes**

Mitochondria were isolated from UC-MSCs by differential centrifugation. $1 \times 10^7$ cells were harvested and washed with PBS. The cells were resuspended in 1 mL of pre-chilled extraction buffer (210 mM mannitol, 70 mM sucrose, 5 mM HEPES, 1 mM KCl, 1 mM EGTA, and 5 mg/mL BSA, pH 7.2). The mechanical homogenization was performed with a glass cell homogenizer on ice. The homogenate was centrifuged at 600*g* for 10 min at 4°C. The supernatant was centrifuged at 8,000*g* for 15 min at 4°C to collect mitochondria. The mitochondria were washed with 300 μL of pre-chilled respiration buffer (225 mM mannitol, 75 mM sucrose, 10 mM KCl, 10 mM Tris-HCl, and 5 mM $KH_2PO_4$, pH 7.2). The mitochondrial pellet was resuspended in the respiration buffer at a concentration of 1~5 mg/mL (total protein concentration) [13].

For microinjection of GV oocytes, 5~7 pL of mitochondrial suspension or respiration buffer (control) was injected into each GV oocyte using the micromanipulator (Narishige, Tokyo, Japan) equipped with FemtoJet 4i microinjector (Eppendorf, Hamburg, Germany). The injected oocytes were transferred to MEM-alpha with 1 mg/mL BSA in a humidified incubator at 37°C with 5% $CO_2$ for 1 h to repair mechanical damages. Then, the oocytes and their cumulus cells were suspended in a 200-μL tube and spun at 3,700*g* for 1 min. The tube was rotated and spun for 1 min. The reconstructed COCs were subjected to IVM and IVF.

**Granulosa cell culture and conditioned media preparation**

Female mice (6~8 weeks old) were intraperitoneally injected with 10 IU PMSG and sacrificed 46~48 h later. Ovaries were collected in sterilized PBS containing 0.5 IU/mL heparin and washed several times. Follicles were punctured with a sterile needle to release granulosa cells. The cells were washed twice by centrifugation at 600*g* for 5 min. Then, the cells were cultured in MEM-alpha supplemented with 50 mIU/mL follicle-stimulating hormone (FSH), 1% ITS, 10% FBS in a humidified incubator at 37°C with 5% $CO_2$. After 24 h, unattached cells were washed away. The culture media were collected at the first passage with 80% confluency. The media were centrifuged at 1,000*g* for 5 min and the supernatant media were stored at −20°C to use as granulosa cell-conditioned media.

**Induction of UC-MSCs into granulosa cells (iGCs)**

Transwell system (CLS3450, Corning, Sigma-Aldrich) was used for the induction of UC-MSCs into granulosa cells. Differentiation media were MEM-alpha supplemented with 20% granulosa cell-conditioned media, 10% FBS, 0.8% L-glutamine, 1% ITS. UC-MSCs were inoculated in 6-well plates and cultured for 24 h. Granulosa cells were seeded in the upper chamber and cultured separately for 24 h. The chamber was inserted into the well containing UC-MSCs and the media were replaced with differentiation media. UC-MSCs and granulosa cells were cocultured for 14 days. The media were refreshed every 2 days.

**Construction of iGC-oocyte complexes**

Aggregation of GV oocytes and iGCs was performed according to a previous paper [29] with some modifications. GV oocytes were incubated for about 20~30 s in prewarmed acidic Tyrode's solution (T1788, Sigma-Aldrich) to weaken the zona pellucida rather than remove it.

They were washed in HEPES-MEM plus 1 mg/mL BSA. The zona-weakened GV oocytes and individual or small clumps of iGCs were tenderly mixed by repeated pipetting. 10~15 oocytes and iGCs were deposited into the base of a 200-μL tube containing 100 μL PBS. The tube was spun at 3,700*g* for 1 min and then rotated and spun for 1 min. The tube was cut near the base and the pellet was carefully scooped out. The pellet was cultured in GDF9-containing media (MEM-alpha supplemented with 100 ng/mL GDF9, 10 mIU/mL FSH, 10 nM estradiol, 3 mg/mL BSA) in a humidified incubator at 37°C with 5% $CO_2$ for 3 days. Then, iGC-oocyte complexes were subjected to *in vitro* maturation (IVM) and *in vitro* fertilization (IVF). A subset of zona-intact GV oocytes and their cumulus cells were reconstructed into COCs and used as control.

For observation of mitochondria transport into zona-weakened GV oocytes via transzonal filopodia, we used mitochondria-labeled cells (see below: **UC-MSCs transfection**) to construct iGC-oocyte complexes. A subset of zona-intact GV oocytes were also aggregated with iGCs into complexes and used as control. After 3 days of incubation, the mass of cells was gently teased apart and some of the iGCs were removed by repeated aspiration for clearer visualization of mitochondria transport through transzonal filopodia. The nuclei of iGC-oocyte complexes were stained with Hoechst 33342. The images for the complexes were captured by using the Z-stack mode under a Nikon A1+ laser confocal microscope.

To examine whether the zona pellucida hampered mitochondria transport, the zona was punched with a 7 μm diameter glass capillary (World Precision Instruments, Sarasota, FL). The aggregation and imaging were performed as above.

**UC-MSCs transfection**

UC-MSCs were inoculated in a 60 mm dish with 4 mL DMEM-LG plus 10% FBS, cultured to 70~80% confluence. Cells were digested with 0.25% trypsin and washed twice with Opti-MEM. Then cells were resuspended and adjusted to $5 \times 10^6$ cells/mL with Opti-MEM. 20 μg plasmid (tdTurboRFP-Mito-7 was a gift from Michael Davidson. Addgene plasmid# 58062; http://n2t.net/addgene:58062; RRID: Addgene_58062) was added to the cells. Then, 200 μL of this mixture was transferred to a 4 mm-Gap cuvette. Electroporation was done using a BTX ECM 2001 electroporator (Harvard Apparatus, Holliston, MA) with the parameter of 500 V, 1 ms, 3 pulses. After 10 min of equilibration, cells were inoculated in a 90 mm dish and cultured for 24 h. Then, the media were replaced with DMEM-LG containing 10% FBS and 800 μg/mL G418. After 7 days, the G418 concentration was reduced to 200 μg/mL. The G418-containing media were refreshed every 3 days until monoclonal colonies emerged. The RFP-positive colonies were picked out and expanded for the construction of iGC-oocyte complexes.

**Electron microscopy**

The oocytes or cell pellets were fixed overnight in 2.5% glutaraldehyde in PBS at 4°C. Low melting point (LMP) agarose was prepared in a 3 $mm^3$ cubic mold. After solidification, a 1 mm diameter screwdriver head was passed back and forth on the flame, stopped for a short while, and then pressed down vertically into the LMP agarose block to form a 1 $mm^3$ microwell. The oocytes or cell pellets were transferred into the microwell and covered with 37°C LMP agarose. This agarose block was then fixed, dehydrated, embedded, and ultramicrotomed. The ultrathin sections were stained with uranium acetate and lead citrate and observed under a transmission electron microscope (HT7700, Hitachi High-Tech, Minato-ku,

Tokyo, Japan). To quantify the proportion of abnormal mitochondria in the GV oocyte, the number of normal mitochondria and abnormal mitochondria (with ultrastructural features including the mitochondrial vacuole, a narrowed inter-membrane space, and myelin figures) was counted in different ooplasm regions of several oocytes.

**IVM, IVF, embryo culture, and embryo transfer**

Every 10 reconstructed COCs or iGC-oocyte complexes were cultured in a 50 μL droplet of MEM-alpha supplemented with 0.2 IU/mL FSH, 1.5 IU/ml human chorionic gonadotropin, 10 ng/mL EGF, 1% ITS, 3 mg/mL BSA for 16 h in a humidified incubator at 37°C with 5% $CO_2$.

For IVF, sperm were collected from cauda epididymides of a 12-week-old male and incubated in 1 mL of HTF (human tubal fluid) medium for 1 h in a humidified incubator at 37°C with 5% $CO_2$. After capacitation, spermatozoa were added to a 300 μL HTF droplet containing mature reconstructed COCs or iGC-oocyte complexes at a final concentration of $1 \times 10^6$ spermatozoa/mL. After 4~6 h of incubation, inseminated oocytes were thoroughly washed to remove adhered spermatozoa and cumulus cells by repeated pipetting.

Every 8~12 presumptive zygotes were cultured in a 40 μL droplet of KSOM-AA (potassium simplex optimized medium with amino acids) containing 8 mg/mL BSA at 37°C under a humidified atmosphere of 5% $CO_2$ in air (0 h). The development to 2-cell or blastocyst stage was examined at 24 h or 84 h, respectively.

For embryo transfer, pseudopregnant recipients were generated by mating females with vasectomized males. *In vitro* cultured 2-cell embryos were transferred into oviducts (4~7 embryos/oviduct; 6~8 microinjected embryos/oviduct) of 0.5 dpc (days post-coitus) pseudopregnant recipients.

**ATP content measurement**

The ATP content in oocytes was measured using Enhanced ATP Assay Kit (S0027, Beyotime) according to the user's manual. Briefly, 100 μL of ATP Assay Working Solution was added into each well of a 96-well white plate and placed at room temperature for 3~5 min. During this period, oocytes were deposited individually in 20 μL of ATP Releasing Solution on ice. Then, they were added to ATP Assay Working Solution and measured using a SpectraMax iD3 Microplate Reader (Molecular Devices, San Jose, CA). Standard curve was established by measuring serial dilutions of ATP Standard (0, 0.02, 0.05, 0.1, 0.2, 0.4, 0.6, 0.8, 1.0, 2.0, 4.0 pmol of ATP). The ATP content was defined according to the standard curve.

**Reactive oxygen species assay**

The intracellular ROS levels in oocytes were detected by using a Reactive Oxygen Species Assay Kit (S0033, Beyotime). The oocytes were incubated in KSOM-AA containing 4 μM 2′,7′-dichlorodihydrofluorescein diacetate (DCFH-DA) for 30 min. Then they were washed 3 times in PBS. Immediately, the ROS levels in oocytes were detected at 488 nm under an A1+ laser confocal microscope. Images were acquired by using the same confocal microscope settings. For quantitative analysis of ROS levels in the embryos, fluorescence images were analyzed by using ImageJ (×1.38) software (https://imagej.nih.gov/ij/) according to the user guide. The image background was subtracted, and then the pixel value of fluorescence was measured using the region of interest (ROI) function.

**Criteria of spindle scoring**

Normal spindles in MII oocytes showed a symmetrically barrel-shaped morphology with condensed chromosomes positioned linearly on the equatorial plate. Abnormal spindles exhibited the following characteristics: a reduction in the number of microtubules; shortened, asymmetric spindles with disordered fibers; dissolved or missing spindles; disordered chromosomes clutter distribution between the spindle fibers; the loose arrangement of chromosomes; nearly normal microtubule structure yet with misaligned and detached chromosomes was also defined as abnormal.

**Apoptosis detection**

The number of apoptotic cells in cultured blastocysts was determined using a DeadEnd Fluorometric TUNEL System (G3250, Promega (Beijing) Ltd., Beijing, China). Between two steps, blastocysts were washed three times in PBS/PVP for 5 min, unless otherwise stated. Blastocysts were fixed in 4% paraformaldehyde in PBS for 20 min and permeabilized in 0.2% Triton X-100 in PBS for 5 min. Then they were equilibrated in equilibration buffer for 8 min and then incubated with FITC-conjugated dUTP and terminal deoxynucleotidyl transferase in equilibration buffer at 37°C for 60 min in the dark. The tailing reaction was terminated in 2× SSC for 15 min. Next, blastocysts were blocked in blocking solution for 1 h. Whereafter, they were immediately incubated in 1: 100 dilutions of mouse anti-CDX2 antibody (AM392, RRID: AB_2650531, BioGenex, Fremont, CA) in blocking solution overnight at 4°C. Antibodies that bound to blastocysts were probed with Cy3-conjugated donkey anti-mouse secondary antibody (1: 800 dilutions, 715-165-151, RRID: AB_2315777, Jackson ImmunoResearch) in blocking solution for 20 min. The nuclei were counterstained with DAPI. Blastocysts were observed under an A1+ laser confocal microscope. The apoptotic index indicated the incidence of apoptotic cells in blastocysts and was calculated via the formula: (apoptotic cell number/total cell number) × 100.

**Experimental design**

The umbilical cord was collected from the female pup. Then, its UC-MSCs were isolated and cryopreserved.

After the female aged (10~12 months old) with decreased fertility, its GV oocytes were collected and treated to weaken the zona pellucida. Its autologous UC-MSCs were induced into granulosa cells (iGCs). An aged GV oocyte was aggregated with its autologous iGCs into a complex. Then, the iGC-oocyte complexes were cultured in GDF9-containing media for 3 days.

The iGC-oocyte complexes were subjected to *in vitro* maturation and fertilization. Presumptive zygotes were cultured for 24 h, the cleaved 2-cell embryos were selected for embryo transfer.

The proposed strategy for oocyte rejuvenation was illustrated in Figure 6.

**Statistical analysis**

Each experiment was independently performed at least three times. The exact values of sample number (n) were shown in the figures or legends. The data are presented as mean ± SEM unless otherwise indicated. Differences between experimental groups were analyzed by Student's *t*-test or one-way ANOVA followed by a Holm-Sidak test using SigmaStat3.5 (Systat Software Inc., Palo Alto, CA). Values of $P < 0.05$ were considered significantly different. The exact values for both significant and non-significant $P$ values were provided in the figures or legends.

**Table S1. Primers used in RT-PCR**

| Gene | Primers | Product (bp) |
|---|---|---|
| *Gapdh* | GTGTTCCTACCCCCAATGTGT | 248 |
| | ATTGTCATACCAGGAAATGAGCTT | |
| *Nanog* | GCAGAAGTACCTCAGCCTCCA | 224 |
| | ATGCGTTCACCAGATAGCCCT | |
| *Sox2* | ACCGATGCACCGCTACGACG | 200 |
| | TGGAGTGGGAGGAAGAGGTAACCA | |
| *Tert* | CTCCTGTCGGTCTTGCGGTTG | 166 |
| | GGTTCTTCCTAACACGCTGGTCA | |
| *Rex1* | CCAAGTGTTGTCCCCAAATAC | 213 |
| | GTCTTGCTTTAGGGTCAGTCTG | |
| *Oct4* | GCAGATCACTCACATCGCCAATC | 133 |
| | TCCCTGTAGCCTCATACTCTTCTCGT | |
| *Amh* | CTCTGATTCCCGCTGTTTCACG | 334 |
| | TAATAGGGGTTCCTCCCAGTCGA | |
| *Amhr2* | GCCAGAATGTGCTCATTCGG | 493 |
| | CACTCAGCTGTCAGCCGTGC | |
| *Fshr* | TAGGATTGAAAAGGCTAACAATC | 210 |
| | AAGCTCAGTCCCATGAAGGA | |
| *Cyp19a1* | GGCATCATATTTAACAACAACCCG | 171 |
| | CACATCCACGTAGCCCGAGG | |
| *Foxl2* | GCCTGCGAGGACATGTTCGAG | 210 |
| | GTTGTTGAGGAACCCCGATTGC | |

**Table S2. Primary antibodies used in immunofluorescence**

| Antibodies (dilutions) | Source | Catalog No. | RRID |
|---|---|---|---|
| rabbit anti-AMHR2 (1: 100) | Thermo Fisher | PA5-112901 | AB_2867635 |
| goat anti-AMH (1: 100) | R&D System | AF1446 | AB_2226486 |
| rabbit anti-FSHR (1: 200) | BIOSS | bs-0895R | AB_10855099 |
| goat anti-FOXL2 (1: 500) | R&D System | NB100-1277 | AB_2106187 |
| rabbit anti-OCT4 (1: 400) | Cell Signaling Technology | 2840 | AB_2167691 |
| rabbit anti-SOX2 (1: 400) | Cell Signaling Technology | 23064 | AB_2714146 |
| rabbit anti-NANOG (1: 1,000) | Cell Signaling Technology | 8822 | AB_11217637 |
| rabbit anti-KLF4 (1: 500) | Thermo Fisher | PA5-27441 | AB_2544917 |
| mouse anti-α-Tubulin (1: 200) | Thermo Fisher | 62204 | AB_1965960 |

**Table S3. Secondary antibodies used in immunofluorescence**

| Secondary Antibodies | Source | Catalog No. | RRID |
|---|---|---|---|
| donkey anti-rabbit IgG, Cy2 conjugated | Jackson ImmunoResearch | 711-225-152 | AB_2340612 |
| donkey anti-rabbit IgG, Cy3 conjugated | Jackson ImmunoResearch | 711-165-152 | AB_2307443 |
| donkey anti-goat IgG, Cy2 conjugated | Jackson ImmunoResearch | 705-225-147 | AB_2307341 |
| donkey anti-goat IgG, Cy3 conjugated | Jackson ImmunoResearch | 705-165-147 | AB_2307351 |
| donkey anti-mouse IgG, Cy2 conjugated | Jackson ImmunoResearch | 715-225-151 | AB_2340827 |

**Table. S4. Antibodies used in flow cytometry**

| FACS Antibodies | Source | Catalog No. | RRID |
|---|---|---|---|
| rat anti-CD19, PE-conjugated | eBioscience (Thermo Fisher) | 12-0193-82 | AB_657659 |
| rat anti-CD90, PE-conjugated | eBioscience (Thermo Fisher) | 12-0902-82 | AB_465776 |
| rat anti-CD105, PE-conjugated | eBioscience (Thermo Fisher) | 12-1051-82 | AB_657524 |
| rat anti-CD45, FITC-conjugated | eBioscience (Thermo Fisher) | 11-0451-82 | AB_465050 |
| rat IgG2a, PE-conjugated | eBioscience (Thermo Fisher) | 12-4321-80 | AB_1834380 |
| rat IgG2b, FITC-conjugated | eBioscience (Thermo Fisher) | 11-4031-82 | AB_470004 |
